# CURLYLEAF is a key modulator of apoplast water status in Arabidopsis leaf

**DOI:** 10.1101/2022.09.23.509124

**Authors:** Jingni Wu, Jinyu Zhang, Xiao Mei, Luhuan Ye, Tao Chen, Yiping Wang, Menghui Liu, Yijing Zhang, Xiu-Fang Xin

## Abstract

The apoplast of plant leaf, the intercellular space between mesophyll cells, is normally largely filled with air with a minimal amount of water in it, which is essential for key physiological processes such as gas exchange to occur. Interestingly, phytopathogens exploit virulence factors to induce a water-rich environment, known as “water soaking”, in the apoplast of the infected leaf tissue to promote disease. We propose that plants evolved a “water soaking” pathway, which normally keeps a “minimized and balanced” water level in the leaf apoplast for plant growth but is disturbed by microbial pathogens to facilitate infection. Investigation of the “water soaking” pathway and leaf water control mechanisms is a fundamental, yet previously-overlooked, aspect of plant physiology. To identify key components in the “water soaking” pathway, we performed a genetic screen to isolate Arabidopsis *severe water soaking* (*sws*) mutants that show leaf water over-accumulation under high air humidity, a condition required for visible water soaking. Here we report the *sws1* mutant, which displays rapid water soaking upon high humidity treatment due to a loss-of-function mutation in *CURLY LEAF (CLF)*, encoding a histone methyl-transferase in the POLYCOMB REPRESSIVE COMPLEX 2 (PRC2). We found that the *sws1 (clf)* mutant exhibits an enhanced abscisic acid (ABA) level and stomatal closure, which are indispensable for its water soaking phenotype and mediated by CLF’s direct regulation of a group of ABA-associated NAC transcription factors, *NAC019/055/072*. Interestingly, the *clf* mutant showed a weakened immunity, which likely also contributes to the water soaking phenotype. In addition, the *clf* plant supports a significantly higher level of *Pseudomonas syringae* pathogen-induced water soaking and bacterial multiplication, in an ABA pathway and *NAC019/055/072*-dependent manner. Collectively, our study probes into a fundamental question in plant biology and demonstrates CLF as a key modulator of leaf water status via epigenetic regulation of ABA pathway and stomatal movement.

## Introduction

Water is essential for virtually all biological processes in living organisms and water availability largely affects plant diversity and microbial community (Lau and Lennon, 2012; Blazewicz et al., 2014; Jonas et al., 2015; Taketani et al., 2017). As sessile organisms, plants evolved mechanisms to sense and respond to environmental water status. For example, plants adapt to water deprivation, drought or flooding conditions through abscisic acid (ABA)- and ethylene-related pathways and reprogram growth (Loreti et al., 2016; Kuromori et al., 2018; Zhao et al., 2017; Zhu, 2016). Plant diseases caused by foliar pathogens are also strongly influenced by water status in the plant and the environment (i.e., air humidity). For instance, *Pseudomonas syringae* and *Xanthomonas gardneri* bacterial pathogens utilize effectors, a class of virulence proteins produced by Gram-negative bacteria and delivered into plant cells through the type III secretion system, to induce water accumulation in the apoplastic space of the infected leaf tissue. This aqueous living environment, also called “water soaking”, is important for pathogen multiplication or egression (Xin et al., 2016; Schwartz et al., 2017). Therefore, foliar pathogens purposely modify the apoplast water availability to cause disease. However, plants normally maintain a “dry” apoplast with a minimal amount of water, which is unarguably important for physiological processes such as gas exchange and photosynthesis to occur. How the apoplastic water status is regulated in the plant leaf is fundamentally-important but poorly-studied at the moment.

The ABA hormone pathway mediates adaptive responses of plants to water deficient or osmotic stress conditions (Chen et al., 2020). ABA is perceived by the pyrabactin resistance (PYR) and PYR1-like (PYL) receptors, which interact with protein phosphatases 2C, and results in the released activity of sucrose non-fermenting 1-related protein kinase 2s (SnRK2s), including SnRK2.2, SnRK2.3 and SnRK2.6 (Cutler et al., 2010; Fàbregas et al., 2020). These SnRK2s in turn activate transcription factors to induce the expression of downstream genes, including *9-cis-epoxycartenoid dioxygenase 3 (NCED3)* for ABA synthesis and other ABA-responsive genes for metabolic and physiological responses (Hewage et al., 2020). SnRK2s, particularly SnRK2.6 (also named Open Stomata 1, OST1), also phosphorylate and activate plasma membrane-located anion efflux channels, namely slow anion channel 1 (SLAC1) and SLAC1-homolog proteins 3 (SLAH3), in the guard cell to trigger stomatal closure and minimize water loss (Hedrich and Geiger, 2017). SLAC1 and SLAH3 are also regulated by calcium-dependent protein kinases (CPKs) (Chen et al., 2020), and both can inhibit the inward-rectifying K+ channel KAT1 and trigger stomatal closure (Zhang et al., 2016). Other stimuli such as reactive oxygen species, darkness and microbe-associated molecules (e.g. flagellin) also trigger stomatal movement (Kwak et al., 2003; Guzel Deger et al., 2015).

Polycomb group (PcG) proteins are epigenetic chromatin modifiers and highly conserved in *Drosophila*, plants and mammals (Köhler and Villar, 2008). PcG proteins form multi-protein complexes, with polycomb repressive complex 1 and 2 (PRC1/2) best studied, to keep the repressed state of target genes and regulate diverse developmental processes. In Arabidopsis, PRC2 deposits the trimethylation mark on the K27 residue of histone 3 (H3K27me3) and the core catalytic subunits are SET domain-containing methyl-transferases, namely Curly Leaf (CLF), Swinger (SWN) and Medea (MEA), which function (partially) redundantly but in distinct processes. PRC1 is comprised of Like Heterochromatin Protein1 (LHP1), which recognizes H3K27me3-modified chromatin (Turck et al., 2007; Zhang et al., 2007), and two groups of RING-domain proteins, AtRING1a/b and AtBMI1a/b/c, which monoubiquitylate histone 2A (Xu and Shen, 2008; Bratzel et al., 2010). PRC1 and PRC2 act together to establish a repressed chromatin status and play pivotal roles in embryogenesis, seed germination, flowering time control and so on (Xiao and Wagner, 2015; Pu and Sung, 2015). Interestingly, many abiotic stress (i.e., cold, dehydration, osmosis)-responsive genes are occupied by H3K27me3 mark and epigenome profiles change under abiotic stress conditions (Liu et al., 2014; Kwon et al., 2009; Liu et al., 2019). CLF/SWN-associated H3K27me3 was shown to repress ABA responses and regulate AB A-induced senescence (Liu et al., 2019). These studies suggest that PRC2 also modulates plant stress responses.

Our study aims to uncover key components involved in controlling water status in Arabidopsis leaf. Through an ethylmethane sulfonate (EMS)-based genetic screen, we identified CLF, the catalytic subunit of PRC2, as a key regulator of apoplast water. The *clf* mutant plant displays a strong spontaneous “water soaking” under high air humidity, and this is tightly linked to activation of ABA pathway and stomatal closure. Blockage of ABA biosynthesis or stomatal movement in the *clf* plant largely compromised the water soaking phenotype. Furthermore, we found a group of ABA-associated NAC transcription factors, *NAC019/055/072*, are directly targeted by CLF and responsible for the water soaking phenotype in the *clf* mutant. Our work identifies a novel function of CLF in modulating leaf water status and illustrates a crucial role of ABA-mediated signaling pathway and stomatal movement in the process.

## Results

### A genetic screen for Arabidopsis *severe water soaking* (*sws*) mutants

To uncover key elements involved in apoplastic water control, an EMS-mutagenized population was generated in the background of *atmin7/fls2/efr/cerk1* (hereinafter *mfec*) mutant, which exhibits enhanced, but not saturated, water soaking compared to Col-0 plants under high humidity (Xin et al., 2016). The M2 population was screened for mutants showing severe water soaking in the leaf under high air humidity (~90-95% relative humidity), a condition required for water to “stay” in the apoplast and preventing water evaporation from stomata so that visible “water soaking” can be seen. These mutants were named *severe water soaking* (*sws*) mutants. Among those, the *sws1* mutant was identified as a strong allele and exhibits water soaking quickly (i.e., 0.5h) after high humidity treatment (Figure 1A).

**Figure 1.**
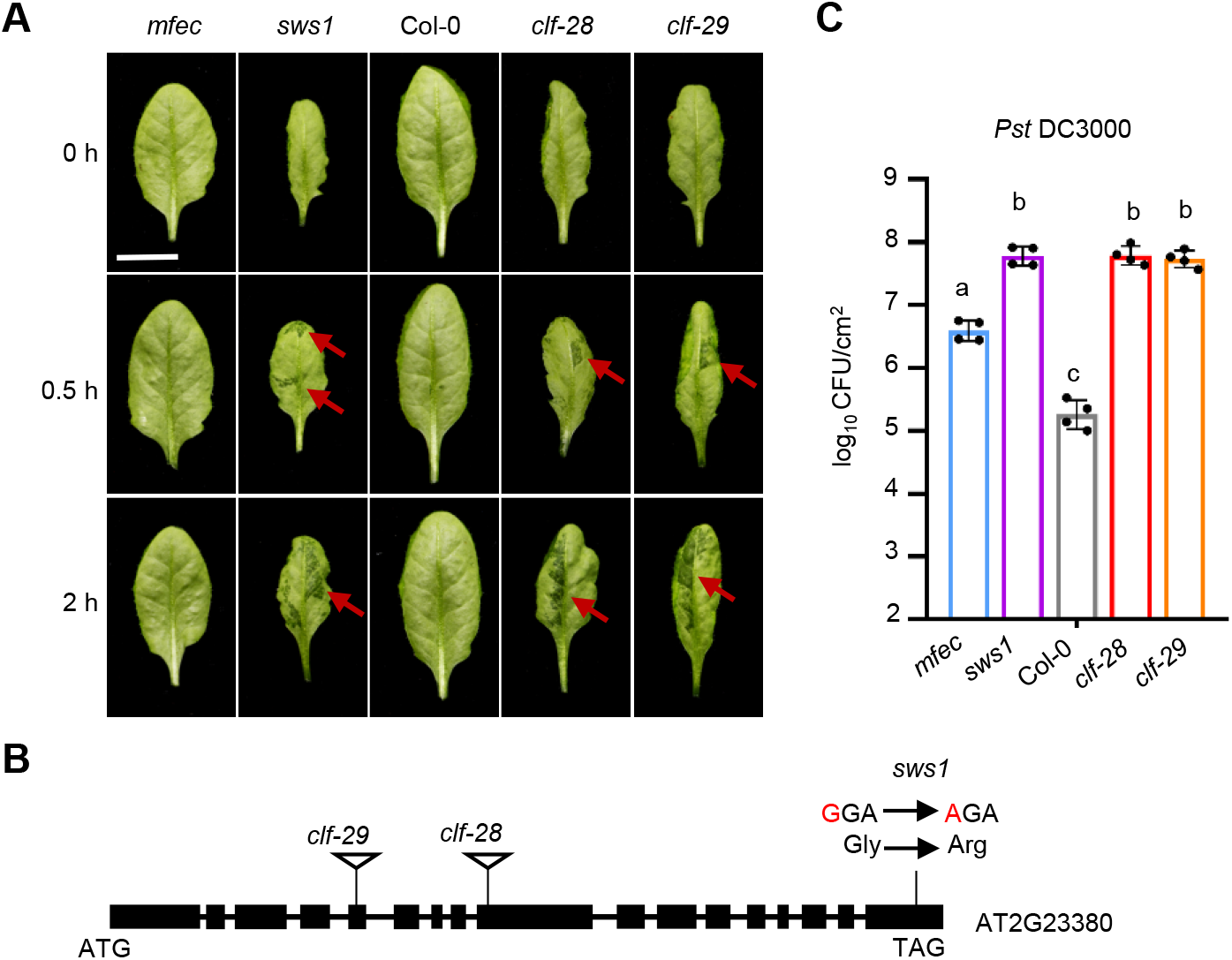
The *severe water soaking1* (*sws1*) mutant exhibits strong water soaking under high humidity and increased susceptibility to *P. syringae* infection. (A) High humidity (~90-95% relative humidity) induced water soaking in *mfec, sws1*, Col-0, *clf-28*, and *clf-29* plant leaves. Scale bar = 1 cm. Red arrows indicate water soaking regions. (B) A schematic diagram indicating the mutated site in *sws1* mutant and the T-DNA insertion position in *clf-28* and *clf-29* lines. (C) Bacterial growth in *mfec, sws1*, Col-0, *clf-28* and *clf-29* leaves. *Pst* DC3000 bacteria were syringe-infiltrated into plant leaves at the dosage of 1 x10^6^ CFU/ml and bacterial population were determined at 0 and 2 days post infiltration (dpi). Data are represented as mean ± SD (n=4). One-way ANOVA with Tukey test (P <0.05) was performed. Different letters indicate statistically significant differences.

To identify the responsible gene locus, we crossed *sws1* with the parent plant. A segregation ratio of 1:3 in water soaking phenotype was observed in the F2 population (Supplemental Figure S1A, B), suggesting that a single recessive mutation causes *sws1* phenotype. We also crossed *sws1* to Col-0 plant and observed a segregation ratio of 1:3 in the water soaking phenotype among F2 plants, suggesting that the *sws1* phenotype is independent of the *mfec* background (Supplemental Figure S1B). Bulked segregant analysis (BSA) coupled with genome sequencing of the F2 population of *sws1* x *mfec* crossed plants was performed and results revealed a G to A nucleotide substitution in the 17th exon of *CURLYLEAF (CLF;* At2g23380) gene, resulting in a Glycine to Arginine substitution (Figure 1B). We then checked another two independent T-DNA insertion lines of *CLF, clf-28* and *clf-29* (Zhou et al., 2017; Bouveret et al., 2006), and water soaking occurred at a comparable level to *sws1* plants in these plants (Figure 1A), suggesting that the mutation in the *sws1* mutant knocked out CLF activity. Notably, another seven independent *sws* lines isolated from the screen also show SNPs in the *CLF* gene (Supplemental Figure S2), suggesting that *CLF* is a mutation hot spot and that our screen is near saturation.

It’s known that water soaking in the plant leaf promotes pathogen infection (Xin et al., 2016). We therefore performed a bacterial growth assay, in which the *P. syringae* pv. *tomato (Pst)* DC3000 bacterium was infiltrated into the leaves of *sws1* (*mfec* background), *clf-28* and *clf-29* (Col-0 background) plants. We found that *Pst* DC3000 grew significantly higher in the *clf* mutants compared to the control plant (Figure 1C), consistent with an important role of apoplast water in bacterial pathogenesis (Xin et al., 2016).

### The aqueous apoplast in the *clf* mutant is mediated by up-regulation of ABA pathway

To understand the physiological basis of the aqueous apoplast phenotype in *clf* plant, we carried out a transcriptome analysis of Col-0 and *clf-28* (hereinafter *clf)* plants at 0 h and 0.5 h after high humidity treatment. In total, 2059 and 2513 differentially expressed genes (DEGs) were detected at 0 h and 0.5 h, respectively, in the *clf* plant (Figure 2A). Gene Ontology analysis revealed that ABA, jasmonic acid (JA) and salicylic acid (SA) pathway genes are differentially regulated (Figure 2B). To clarify which hormone pathway is biologically relevant, we exogenously sprayed methyl jasmonic acid (MeJA), ABA or Benzothiadiazole (BTH, a chemical analog of SA) on Arabidopsis leaf and examined whether water soaking occurs. Intriguingly, spray of ABA, but not MeJA or BTH, leads to water soaking in Col-0 leaves (Figure 2C). Furthermore, this ABA-induced water soaking was abolished in ABA receptor polymutant *pyr/pyl112458* and signaling poly-mutant *snrk2.2/2.3/2.6* (Zhao et al., 2018; Fujii and Zhu, 2009) (Figure 2D). These results suggest that ABA pathway is an important regulator of apoplast water balance in Arabidopsis leaf.

**Figure 2.**
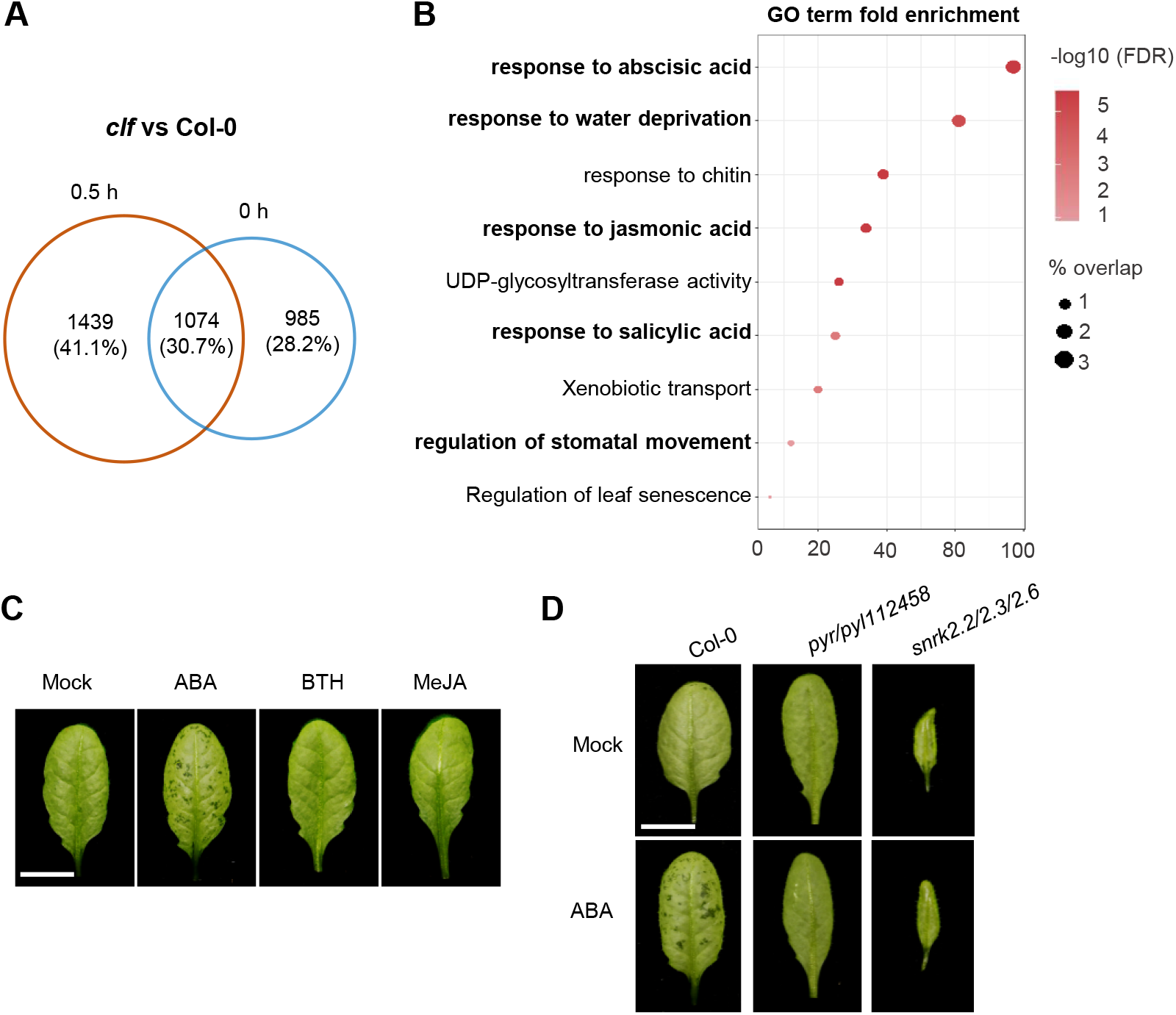
ABA is an important modulator of apoplast water in Arabidopsis leaf. (A) Venn diagram showing the numbers of differentially expressed genes (DEGs) in Col-0 and *clf* plants at 0 and 0.5 h after high humidity treatment. Numbers of DEGs were calculated by pairwise comparison between Col-0 and *clf* (FDR < 0.01; log2 fold change > 1 or < −1). (B) Gene ontology (GO) analysis of all DEGs at 0.5 h after high humidity in the *clf* plant. The X-axis indicates the number of queried genes and Y-axis indicates the GO terms. Size of plotted circles indicates the percentage overlap within input genes. The filled color is scaled to −log10 (FDR). (C) Exogenous application of ABA leads to water soaking in Arabidopsis leaf. ABA (50 μM), BTH (50 μM) or MeJA (50 μM) were sprayed onto Col-0 leaves and plants were covered with transparent plastic dome to keep high humidity. Pictures were taken 8 h later. (D) ABA-induced water soaking was dramatically reduced in *pyr/pyl112458* and *snrk2.2/2.3/2.6* plants. ABA (50 μM) was sprayed onto Arabidopsis leaves and pictures were taken 8 h later. Scale bar = 1 cm.

We then measured ABA level in Col-0 and *clf* plants and found that, while high humidity reduced ABA levels in both plants, the *clf* mutant accumulated a significantly higher level of ABA than Col-0 plant before and after high humidity treatment (Figure 3A). To examine if the altered ABA pathway is responsible for water soaking phenotype, we crossed the *clf* mutant to two ABA mutants, *aba2*, which lacks a key ABA biosynthetic enzyme (González-Guzmán et al., 2002), and *ost1(snrk2.6)*, which lacks a central kinase mediating downstream ABA responses and stomatal closure (Yoshida et al., 2006). The water soaking phenotype was examined in the *clf/aba2* and *clf/ost1* double mutants. We found that water soaking was abolished in the *clf/aba2* plant and largely delayed in the *clf/ost1* plant (Figure 3B), suggesting that the up-regulated ABA pathway mediates the water soaking phenotype in the *clf* plant.

**Figure 3.**
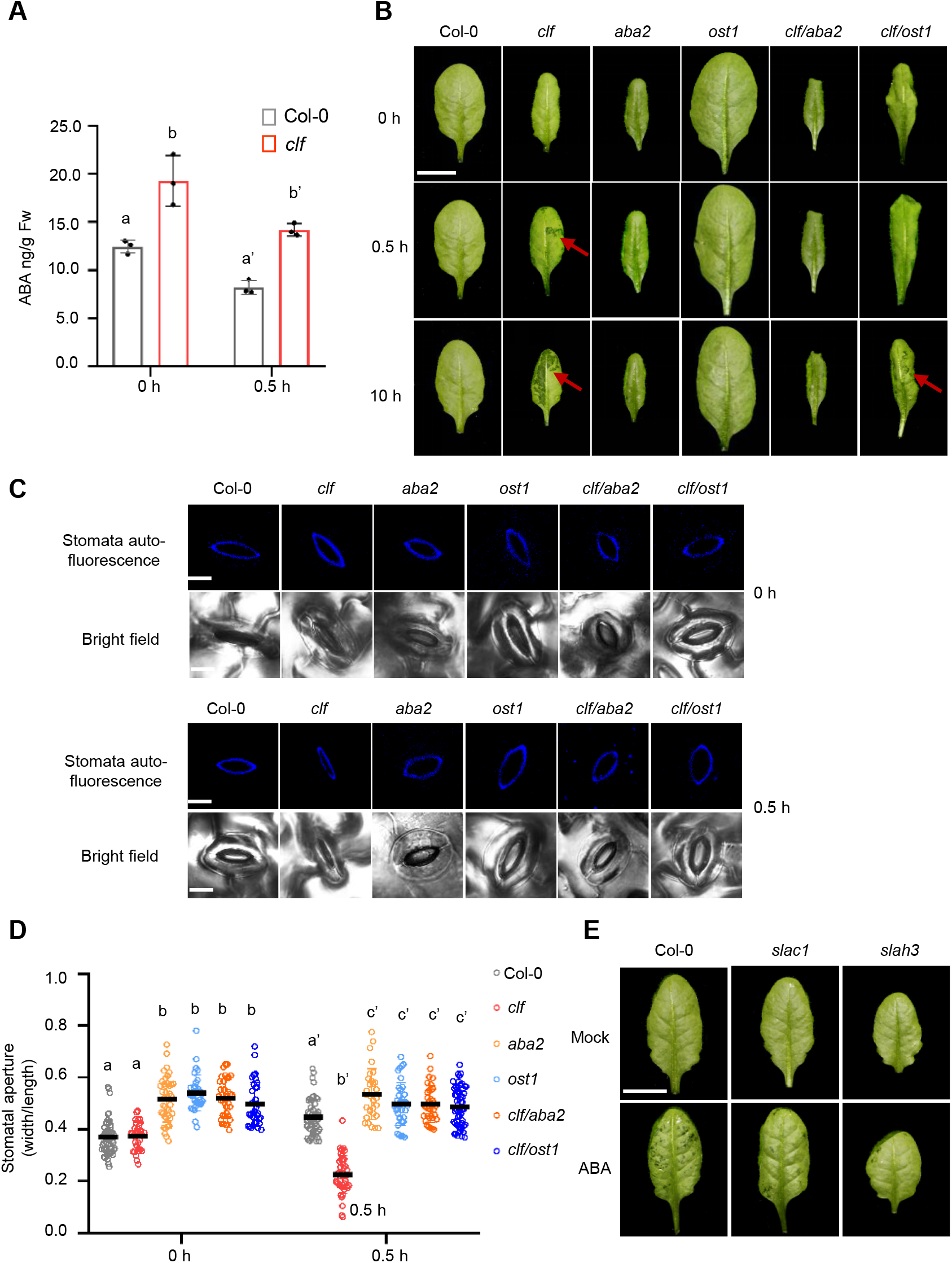
ABA signaling pathway and stomatal closure mediate the water soaking phenotype in the *clf* plant. (A) ABA levels in Col-0 and *clf* plants before and after high humidity. Data are represented as mean ± SD. Different letters indicate significant differences, as determined by two-way ANOVA with Tukey test (P <0.05). (B) Water soaking in Col-0, *clf, aba2, ost1, clf/aba2* and *clf/ost1* plants before and after high humidity treatment. Bar = 1 cm. Red arrows indicate water soaking regions. (C, D) Stomatal aperture in Col-0, *clf, aba2, ost1, clf/aba2* and *clf/ost1* leaves at 0 and 0.5 h after high humidity. Representative stomatal images collected on confocal microscope are shown (bar =10 μm; C) and stomatal aperture (width/length ratio) was calculated (>50 stomata; D). The colored dots represent stomatal aperture of individual stomata. The black line represents the mean. Different letters indicate significant differences, as determined by two-way ANOVA with Tukey test (P <0.05). (E) Water soaking in Col-0, *slac1* and *slah3* plants before or after spray of ABA (50 μM) and being kept under high humidity for 8 h. Bar = 1 cm.

### ABA-associated stomatal movement is essential for aqueous apoplast

Stomatal closure is a key physiological output of ABA signaling transduction (Hsu et al., 2021). As OST1/SnRK2.6 is a key regulator of stomatal closure, the severely-compromised water soaking in the *clf/ost1* plant suggests that stomatal movement is important for water soaking. We then measured the stomatal aperture in the *clf*, *aba2*, *ost1*, *clf/aba2* and *clf/ost1* plant leaves before and after high humidity treatment. As shown in Figure 3C, D, while high humidity led to stomatal opening in Col-0 plants, we observed a striking stomatal closure response in the *clf* leaves 0.5 h after high humidity. Importantly, this stomatal closure did not occur in the *clf/aba2* or *clf/ost1* plants (Figure 3C, D). We further checked the water soaking level in the Arabidopsis *slac1* and *slah3* mutants, which are mutated in two key anion channels in the guard cell and defective in stomatal closure (Zhang et al., 2016), and found that ABA-induced water soaking is much weaker in these two mutants compared to that in Col-0 (Figure 3E). Taken together, these results support a crucial role of stomatal movement in regulating apoplastic water status.

### CLF directly targets ABA-associated NAC transcription factors for regulating apoplast water

We next searched for direct targets of CLF that are involved in water regulation. *CLF* encodes a H3K27 methyltransferase and deposits H3K27me3 on chromatin (Goodrich et al., 1997), we thus profiled H3K27me3 loci by chromatin immunoprecipitation coupled with sequencing (ChIP-seq), in wild type and *clf* plants before and after high humidity treatment. Since H3K27me3 is a transcriptional repression marker, genes directly targeted by CLF are expected to show a higher expression level in the *clf* plant. Hierarchical clustering analysis of differentially expressed genes in the *clf* plant under our experimental conditions was performed and revealed five gene clusters (Figure 4A; Supplemental Table S1). Among them, cluster IV represents highly upregulated genes in the *clf* plant (more likely to be direct targets of CLF) at 0.5 h under high humidity, when water soaking was observed, and, importantly, shows the highest percentage of H3K27me3-regulated genes from our ChIP-seq experiment (Figure 4A, right panel). Furthermore, GO analysis of the five clusters showed that genes related to “response to water deprivation” are enriched in cluster IV (Supplemental Figure S3A), supporting that CLF directly targets ABA-associated components to regulate apoplast water.

**Figure 4.**
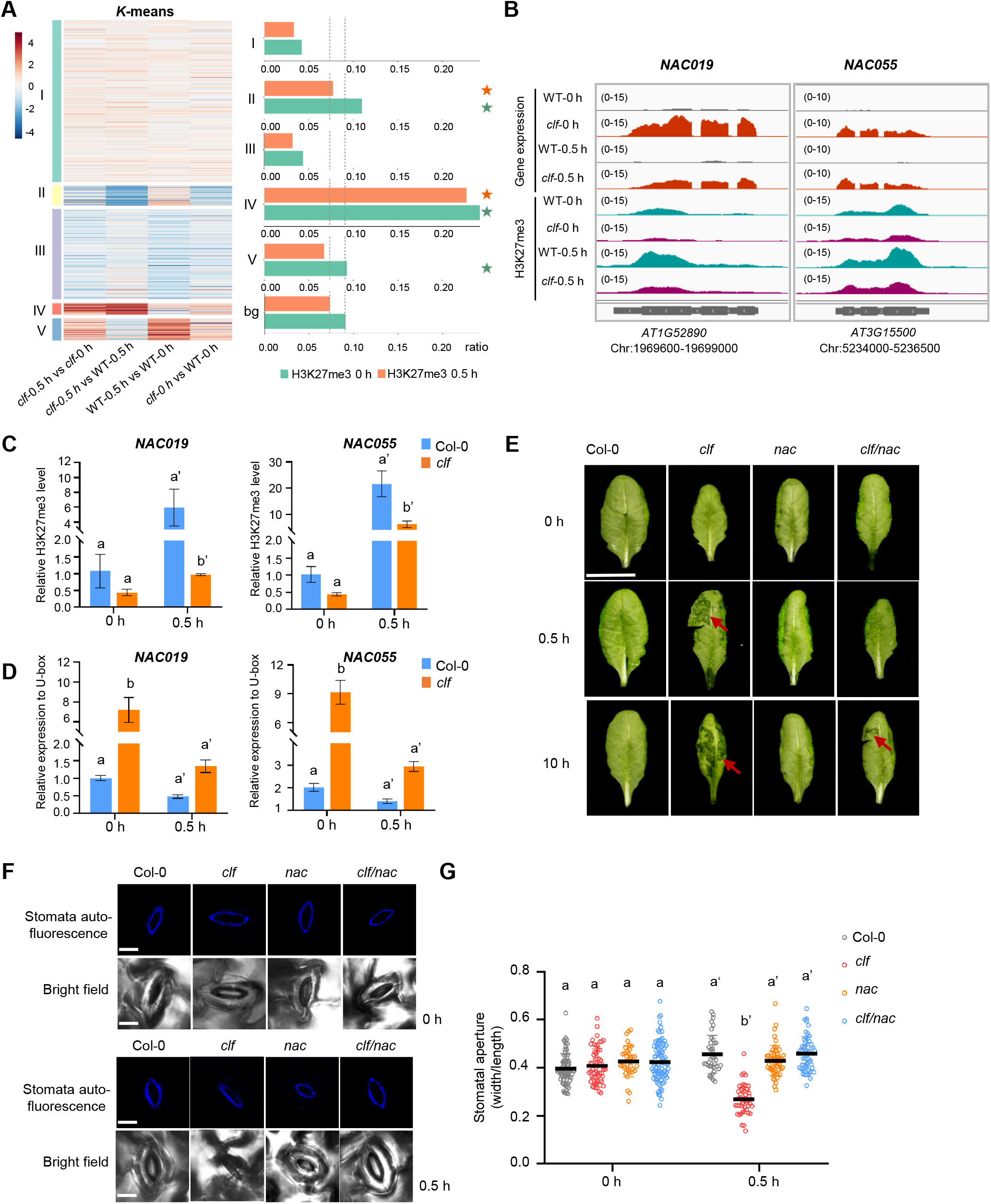
ABA-associated NAC019/055/072 transcription factors are directly targeted by CLF and mediate apoplast water regulation. (A) Hierarchical clustering analysis of the DEGs in Col-0 and *clf* plants at 0 and 0.5 h after high humidity treatment from the RNAseq experiment (FDR < 0.01; log2 fold change<-1 or >1; on the left). The percentage of genes in each cluster showing CLF/H3K27me3 regulation (i.e., differential H3K27me3 level in the *clf* mutant plant from our ChIPseq experiment) are shown on the right. bg, background. (B) Transcription and H3K27me3 levels of *NAC019* and *NAC055* genes in Col-0 and *clf* plants before (0 h) and after (0.5 h) high humidity treatment from RNAseq and ChIPseq experiments. (C) Confirmation of H3K27me3 levels at *NAC019* and *NAC055* genes by ChIP-qPCR. Bars represent means ± SD (n=3). Different letters indicate significant differences, as determined by two-way ANOVA with Tukey test (P<0.05). (D) Confirmation of *NAC019* and *NAC055* expression levels by qRT-PCR. Bars represent means ± SD (n=3). Different letters indicate significant differences, as determined by two-way ANOVA with Tukey test (P <0.05). (E) Water soaking in Col-0, *clf, nac*, and *clf/nac* plants. Plants were treated with high humidity and pictures were taken at indicated time points. Bar=1 cm. (F, G) Stomatal dynamics in Col-0, *clf, nac*, and *clf/nac* plants before and after high humidity. Representative images are shown (bar =10 μm; F) and stomatal aperture is calculated (G). Colored dots represent stomatal aperture of individual stomata (n>50). The black line represents the mean. Different letters indicate significant differences, as determined by two-way ANOVA with Tukey test (P<0.05). Experiments were repeated at least three times with similar results.

Our ChIP-seq results showed that 1527 and 1728 genes, at 0 and 0.5 h after high humidity treatment respectively, showed a differential H3K27me3 level in the *clf* plant, with over 80% of them having a lower H3K27me3 level in *clf* compared to Col-0, indicating that these are candidates of CLF’s direct target genes (Supplemental Figure S3B; Supplemental Table S2). Interestingly, only 769 genes are shared between 0 and 0.5 h datasets, suggesting that different humidity alters the CLF-associated H3K27me3 profile (Supplemental Figure S3B; Supplemental Table S2). Importantly, we found that ABA signaling and homeostasis-related genes, including *PYL6, NPF4.1* and *BG2*, as well as two ABA-inducible NAC transcription factors, *NAC019* and *NAC055*, showed a lower H3K27me3 level, accompanied by a higher gene expression level, in the *clf* plant (Figure 4B; Supplemental Figure S4). The NAC019/055 transcription factors are particularly interesting, because they play an important role in positively regulating ABA responses (Takasaki et al., 2015; Jensen et al., 2010; Tran et al., 2004) and were also identified as CLF targets by previous studies (Liu et al., 2019). ChIP-qPCR and RT-qPCR were performed to confirm the H3K27me3 and transcript levels of *NAC019* and *NAC055* genes in our experimental conditions (Figure 4C, D). To examine if NACs are important for regulating water soaking, we crossed the *clf* plant to the *nac019/nac055/nac072* (hereafter *nac*) mutant, in which three functionally-redundant NACs were knocked out (Takasaki et al., 2015). As shown in Figure 4E, water soaking in the *clf/nac* plant was dramatically delayed compared to the *clf* plant. Furthermore, the rapid stomatal closure phenotype in the *clf* plant was also abolished in the *clf/nac* polymutant (Figure 4F, G). In addition, ABA-induced water soaking was significantly reduced in the *nac* mutant plant (Supplemental Figure S5). These findings suggest that CLF regulates apoplastic water in Arabidopsis leaf, in part, by directly regulating the histone methylation level of NAC transcription factors.

### Plant immunity plays a role in regulating apoplast water

Epigenetic regulation usually exerts a broad role and affects thousands of genes’ transcription. Considering the particularly strong water soaking phenotype in the *clf* plant, we wondered if there are other mis-regulated pathways, in addition to ABA pathway, in the *clf* plant that also contribute to the phenotype. Notably, a large population of SA-responsive genes are differentially regulated in the *clf* plant, compared to Col-0, in our RNAseq experiment (Figure 2B), suggesting that plant immune status is altered in the *clf* plant. To confirm, we tested SA and pattern-triggered immunity (PTI) responses, two primary plant defense pathways (Peng et al., 2021; DeFalco and Zipfel, 2021), in the *clf* mutant. Interestingly, the transcript level of *ICS1*, encoding a rate-limiting enzyme in SA biosynthesis-isochorismate synthase 1, and SA accumulation were significantly reduced in the *clf* plant (Figure 5A, B). In addition, application of flg22, a bacterial flagellin-derived 22-amino acid peptide eliciting PTI (Gómez-Gómez and Boller, 2000; Felix et al., 1999), triggered a much lower ROS production and MAPK phosphorylation in the *clf* plant, compared to Col-0 (Figure 5C, D). These results indicate a weakened immunity in the *clf* plant, suggesting that plant immunity, particularly SA and PTI responses, may negatively affect apoplastic water accumulation. To further test this hypothesis, we examined ABA-induced water soaking in the *ics1* mutant, which is deficient in SA biosynthesis (Wildermuth et al., 2001), and the *bak1-5/bkk1-1/cerk1 (bbc)* triple mutant, which lacks pattern recognition co-receptors and is greatly impaired in PTI responses (Xin et al., 2016). As shown in Figure 5E, stronger water soaking was observed in these two mutants compared with Col-0, suggesting that, in addition to ABA pathway, plant immunity exerts another layer of regulation on apoplast water level. Whether this is through antagonistic crosstalk between ABA and SA/PTI pathways (Moeder et al., 2010; Mine et al., 2017) needs further investigation.

**Figure 5.**
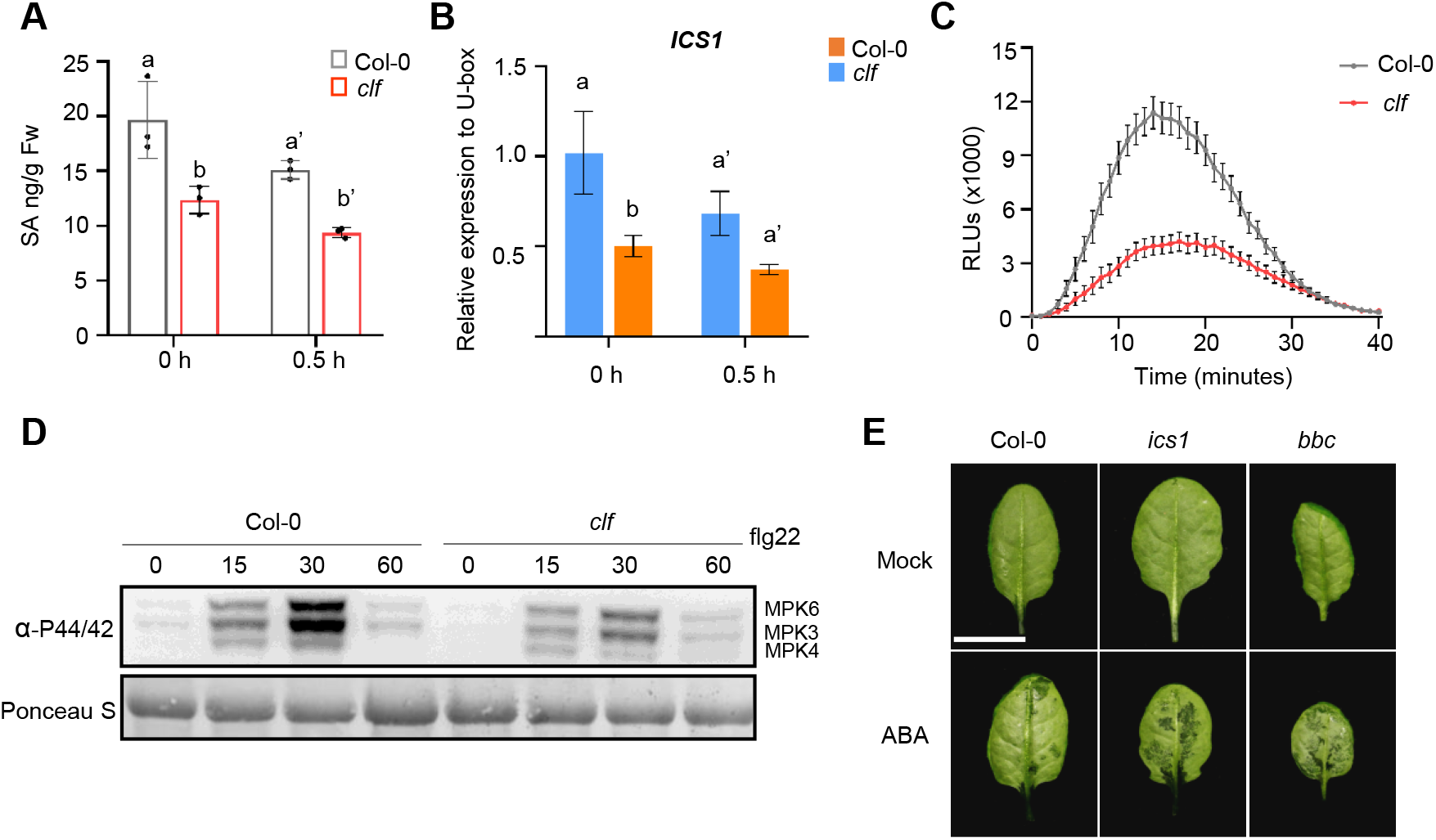
Plant basal immunity seems to play a role in water soaking. (A) SA levels in Col-0 and *clf* plant leaves before and after high humidity treatment, determined by LC/MS/MS. Bars represent means ± SD (n=3). Different letters indicate significant differences, as determined by two-way ANOVA with Tukey test (P <0.05). (B) qRT-PCR analysis of *ICS1* expression in Col-0 and *clf* plants before and after high humidity treatment. Bars represent means ± SD (n=4). Different letters indicate significant differences, as determined by twoway ANOVA with Tukey test (P <0.05). (C) flg22-induced ROS burst in Col-0 and *clf* plants. Leaf disks were treated with 100 nM flg22 and ROS burst was detected by luminol-HRP approach. RLUs, relative luminescence units (n≥20). (D) MAPK phosphorylation in Col-0 and *clf* plants at different time points after 100 nM flg22 treatment. (E) ABA-induced water soaking in Col-0, *ics1* and *bak1-5/bkk1-1/cerk1* (*bbc*) plants. Four-weeks-old plants were sprayed with 25 μM ABA, shifted to high humidity, and pictures were taken 24 h later. Experiments were repeated at least three times with similar results.

### The *clf* mutant exhibits an increased level of *P. syringae-mediated* water soaking and disease susceptibility in an ABA- and NAC-dependent manner

Interestingly, recent studies showed that *P. syringae* bacterial pathogen activates the plant ABA pathway and stomatal closure at a later stage of infection, through type III effectors AvrE and HopM1, to promote water soaking and cause disease (Hu et al., 2022; Roussin-Léveillée et al., 2022). We therefore examined whether the enhanced bacterial susceptibility observed in the *clf* mutant (Fig. 1C) also relies on ABA pathway and stomatal closure. First, we found that *Pst* DC3000 triggered an even stronger stomatal closure and water soaking in the *clf* mutant, compared to Col-0 (Figure 6A-C). Because the *clf* mutant supports a faster bacterial growth, the concentration of inoculum was adjusted so that there were similar bacterial populations in different genotypes when water soaking was observed (24 h post infiltration) (Figure 6C). Importantly, *Pst* DC3000-mediated stomatal closure and water soaking phenotypes were largely abolished in the *clf/aba2* and *clf/ost1* plants, compared to the *clf* mutant (Figure 6A-C). Consistently, while *Pst* DC3000 multiplied to a much higher population in the *clf* plant, this did not occur in the *clf/aba2* or *clf/ost1* plant (Figure 6D).

**Figure 6.**
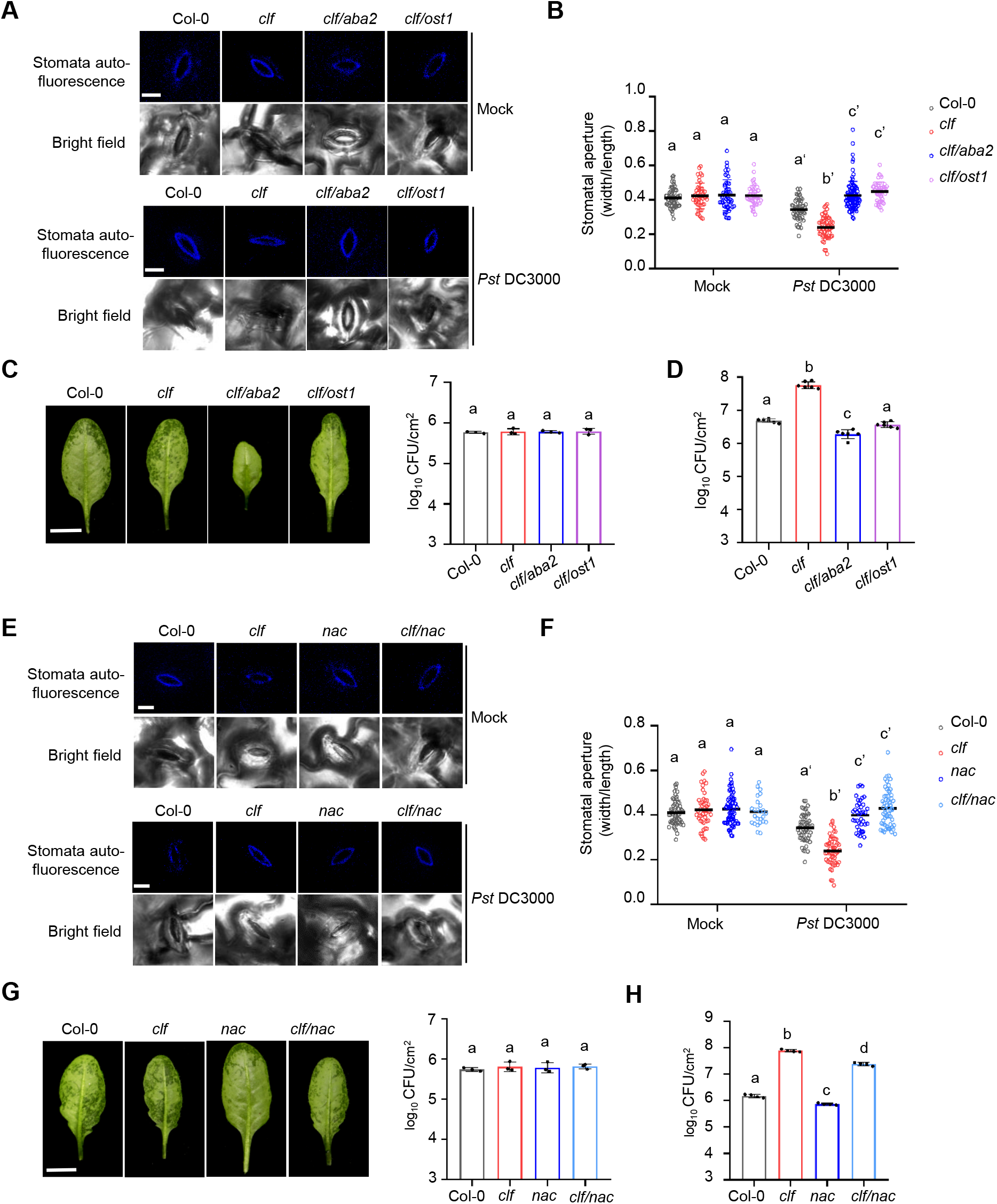
*Pst* DC3000 induced water soaking and disease susceptibility in the *clf, clf/aba2, clf/ost1* and *clf/nac* plants. (A, B) *Pst* DC3000 triggered a stronger stomatal closure in the *clf* mutant, but not the *clf/aba2* or *clf/ost1* plants. Arabidopsis leaves were syringe-infiltrated with Mock (sterile water) or *Pst* DC3000 (at OD_600_=0.1). Stomatal images were collected 4 h after infiltration. Representative images are shown (bar = 10 μm; A) and stomatal aperture is calculated (B). Colored dots represent stomatal aperture of individual stomata (n>40). The black line represents the mean. Different letters indicate significant differences, as determined by two-way ANOVA with Tukey test (P <0.05). (C) *Pst* DC3000 inoculation induced a stronger water soaking in the *clf* plant, but not *clf/aba2* or *clf/ost1* plants. Bacteria were infiltrated into Arabidopsis leaves of Col-0 (1×10^6^ cfu/mL), *clf* (5×10^5^ cfu/mL), *clf/aba2* (4×10^6^ cfu/mL) and *clf/ost1* (2×10^6^ cfu/mL). Pictures were taken 24 h after infiltration (on the left) and bacterial populations were counted (on the right). Bar=1 cm. Different letters indicate significant differences, as determined by one-way ANOVA with Tukey test (P <0.05). (D) The *clf* mutant, but not the *clf/aba2* or *clf/ost1* plants, showed enhanced susceptibility to *Pst* DC3000 inoculation. *Pst* DC3000 bacteria were infiltrated into plant leaves at 1×10^6^ CFU/ml and bacterial titer were determined 2 days post infiltration. Bars represent mean ± SD (n=6). Different letters indicate significant differences, as determined by one-way ANOVA with Tukey test (P <0.05). (E, F) Stomata measurement in Arabidopsis leaves syringe-infiltrated with Mock (sterile water) or *Pst* DC3000 (at OD_600_=0.1). Stomatal images were collected 4 h after infiltration. Representative images are shown (bar = 5 μm; E) and stomatal aperture is calculated (F). Colored dots represent stomatal aperture of individual stomata (n≥40). The black line represents the mean. Different letters indicate significant differences, as determined by two-way ANOVA with Tukey test (P <0.05). (G) *NAC019/055/072* mutations largely compromised the water soaking phenotype in the *clf* plant. Bacteria were infiltrated into Arabidopsis leaves of Col-0 (1×10^6^ cfu/mL), *clf* (5×10^5^ cfu/mL), *nac* (2×10^6^ cfu/mL) and *clf/nac* (1×10^6^ cfu/mL), so that they reached a similar population 24 h after infiltration (on the right). Pictures were taken 24 h after infiltration (on the left). Bar=1 cm. Different letters indicate significant differences, as determined by one-way ANOVA with Tukey test (P <0.05). (H) *NAC019/055/072* mutations largely compromised the enhanced susceptibility in the *clf* plant. *Pst* DC3000 bacteria were infiltrated at 1×10^6^ cfu/mL and bacterial titers were determined 2 days post infiltration. Bars represent means ± SD (n=4). Different letters indicate significant differences, as determined by one-way ANOVA with Tukey test (P <0.05).

Next, we tested if the increased level of bacteria-driven water soaking and stomatal closure in the *clf* mutant also relies on NAC019/055/072 transcription factors. The bacterial infection phenotypes were examined in the *clf/nac* polymutant and, again, we found that the *NAC019/055/072* mutation largely mitigated the stomatal closure and water soaking phenotypes in the *clf* mutant (Figure 6E-G), and, to a lesser degree, the enhanced bacterial growth phenotype (Figure 6H). Taken together, these results suggest that the enhanced water soaking and disease susceptibility during *P. syringae* infection in the *clf* mutant occur through ABA pathway and NAC transcription factors.

## Discussion

Regulation of water status is crucial for various biological processes. Despite the great progress in understanding plant adaptation mechanisms to water deficit conditions in the soil (Kim et al., 2020; Siao et al., 2020; Dinneny, 2019; Ramachandran et al., 2020), how water relations in the leaf tissue is maintained remained elusive. In this study, we genetically isolated the *sws1* mutant that displays abnormal leaf water control and over-accumulated water in the apoplast under high humidity. We show that *SWS1* encodes CURLY LEAF, a key component of the plant PRC2 complex that deposits the H3K27me3 marker on chromatin. We found that CLF normally inhibits ABA pathway by epigenetically silencing ABA-related genes including *NAC019/055/072* transcription factors to keep a dry apoplast, whereas the increased ABA signaling and stomatal closure in the *clf* mutant leads to strong water soaking. Our results also suggest a negative role of plant immunity pathways, particularly SA and PTI pathways, in water accumulation in the leaf apoplast (Fig. 7). Collectively, our study elucidates a new physiological function of CLF, as a key epigenetic regulator, in controlling leaf water balance, likely through a combinatorial regulation of ABA pathway, stomata and plant immunity, shedding light on the elusive basis of water control in the leaf apoplast.

**Figure 7.**
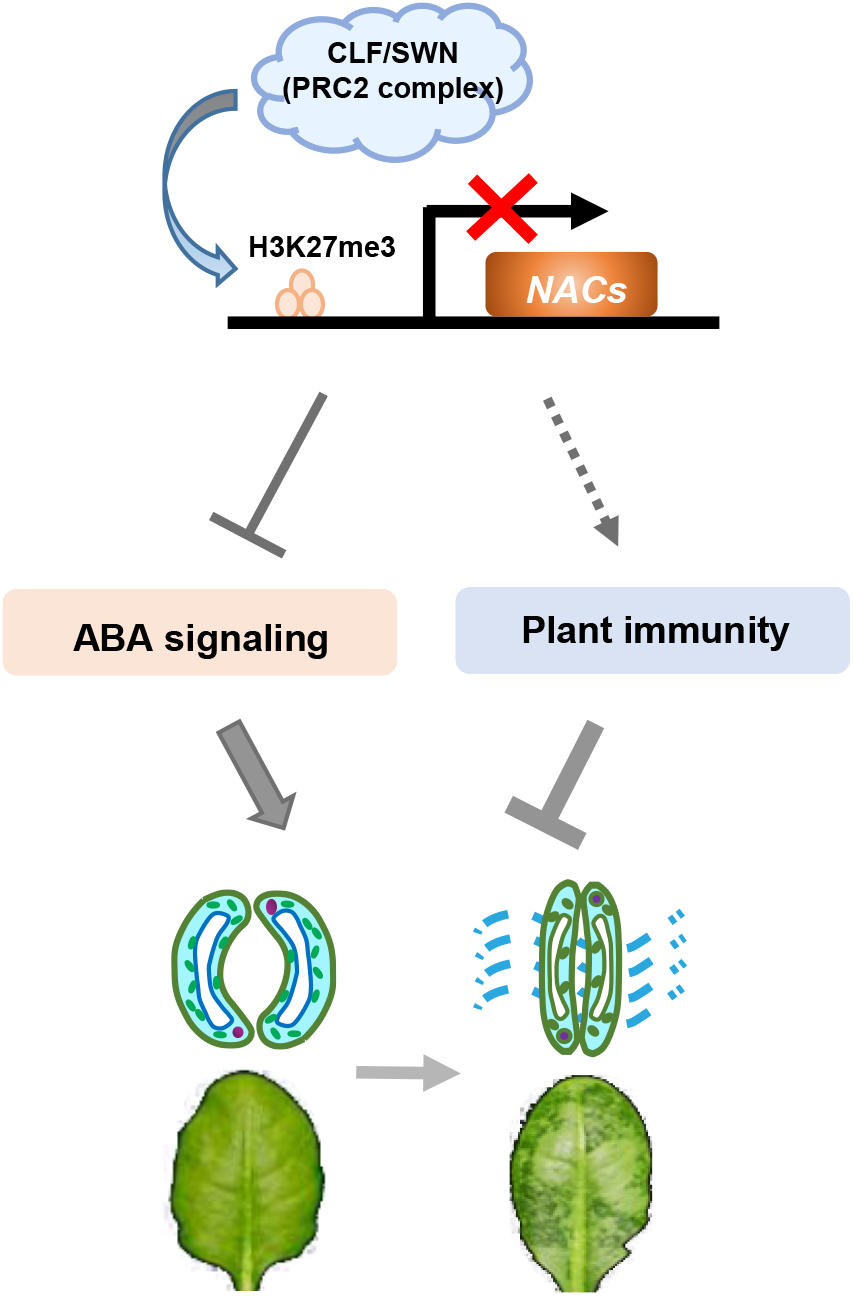
A model describing the findings in this study. The CLF-containing PRC2 complex regulates the H3K27me3 level and chromatin status of ABA pathway genes, including the NAC transcription factor NAC019/055/072, and normally suppresses ABA responses. Absence of CLF (i.e., in the *clf* mutant) results in heightened ABA responses and more closured stomata, leading to water over-accumulation in the leaf apoplast. Plant basal immunity (i.e., SA and PTI) seems to play a negative role in the process.

Previous studies showed that the CLF-containing PRC2 complex epigenetically regulates many ABA-associated genes and influences ABA responses and senescence in Arabidopsis (Liu et al., 2019), which is in agreement with our results. Moreover, our study revealed a striking “humidity-dependent” stomatal aperture dynamics in the *clf* plant, in that a shift to high air humidity quickly (i.e., 0.5 h) triggers a strong stomatal closure in the *clf* plant leaf, presumably leading to water soaking (Figure 3C, D). This phenomenon was not observed in the *clf/aba2* or *clf/ost1* mutants (Figure 3C, D), suggesting that it is an ABA-mediated process. It is possible that CLF exerts a specific, and distinct from mesophyll cells, function in guard cells (e.g., directly regulating the epigenetic and transcriptional status of ion channels for stomatal movement). Detailed mechanisms would need future and cell type-specific investigations. Nonetheless, the striking stomatal closure phenotype in *clf* plant and the fact that it requires key ABA and stomata components supports our main conclusion that CLF modulates leaf water balance through ABA pathway and stomatal movement.

PRC2-mediated H3K27me3 modification is broadly involved in transcriptional gene silencing across plant genome. Interestingly, our study indicates that plant basal immunity may represent another layer of modulation in apoplast water and functions in CLF-regulated leaf water level. This is consistent with the results that water soaking in the *clf* mutant is stronger than that induced by ABA spray treatment (Figure 2C), suggestive of additional pathways, besides ABA, involved in water soaking. We found that *CLF* mutation compromises the accumulation of defense hormone SA and flg22-induced ROS and MAPK phosphorylation (Figure 5A-D), suggesting that CLF directly or indirectly suppresses plant SA and PTI responses. This is in line with a recent study demonstrating an important role of CLF in regulating leaf immunity against fungal pathogens (Singkaravanit-Ogawa et al., 2021) and SA as a negative regulator of pathogen-induced aqueous apoplast (Lajeunesse et al., 2022). It will be interesting to study in the future whether CLF directly regulates certain SA or PTI pathway components through histone methylation, although one cannot rule out the possibility that the lower immunity level in the *clf* plant is due to antagonism between SA/PTI and ABA pathways suggested by previous studies (Cao et al., 2011; Mine et al., 2017). Future investigation of CLF function and other components involved in regulating water relations in the leaf should provide more insights into this fundamental aspect of plant biology.

## Materials and Methods

### Plant materials and growth condition

The *mfec, clf-28, clf-29, nac019/055/072, aba2, ost1, pyr/pyl12458* and *ics1*(also named *sid2*) mutant lines were reported previously (Xin et al., 2016; Zhao et al., 2018; González-Guzmán et al., 2002; Yoshida et al., 2006; Wildermuth et al., 2001). All plants were grown in potting soil in environmentally-controlled growth chamber, with relative humidity set at 60% and temperature at 22 °C and a 12 h light/12 h dark photoperiod. To grow plants on MS plates, seeds were surface sterilized and sown on half-strength MS agar plates containing 0.7% (w/v) agarose. For high humidity treatment, four-to-five weeks old plants were kept at 90-95% relative humidity in an environmentally-controlled growth chamber (MMM group, Germany) for indicated time.

### EMS mutagenesis and isolation of *sws* mutants

The *mfec* seeds were treated with 0.2% ethyl methanesulfornate (EMS) solution for 16 hours. The M1 plants were pooled (six M1 plants were pooled as one family and a total of ~840 families were obtained) and the M2 population were used for screen. M2 seeds were sown in potting soil and plants were grown to three weeks and then covered with a plastic dome to keep high humidity overnight. Plants that displayed severe water soaking the next day were isolated as *sws* mutants.

### BSA analysis and identification of the mutated loci in *sws* mutants

Bulked segregant analysis (BSA) were performed using modified QTL-seq method (Takagi et al., 2013). The read depth information for the homozygous SNPs/InDels in the pools was obtained to calculate the SNP/InDel index. Col-0 and *mfec* genomes were sequenced as reference and for analyzing the read number for the Mut-pool and Wild-pool. Sliding window methods were used to determine the SNP/InDel index. The average of all SNP/InDel indices in each window was used as the SNP/InDel index for that window. A window size range between 2 Mb to 10kb was used as the default settings. The difference between the SNP/InDel index of two pools was calculated as ΔSNP/InDel index. Data analysis was performed on the Majorbio Cloud Platform (www.majorbio.com).

### RNA extraction and real-time RT-qPCR

To analyze gene expression level, four-week-old Arabidopsis plant leave were treated with high air humidity at indicated time points. Three leaves from different plants were collected as one biological replicated and four biological replicates were collected for each treatment. Total mRNA was extracted using TRIzol reagent (Invitrogen, Carlsbad, CA, USA), following the manufacturer’s protocol. Extracted RNA was treated with DNase I (Roche), and two μg of RNA was used to synthesize cDNA by ReverTra Ace® qPCR RT Master Mix with gDNA remover (TOYOBO). Real-time qPCR was carried out with the SYBR Green Realtime PCR master mix (TOYOBO) on a CFX Connect Real Time System (Bio-Rad, Berkeley, CA, USA). The plant *ACTIN2* and *U-box* genes were used as internal standard. Primers used in qPCR are listed in Supplemental Table S3.

### cDNA library generation and RNAseq

For RNAseq experiments, samples were collected as described above. Three leaves from different plants were harvested as one replicate, and four biological replicates were collected for each treatment/time point. Total mRNA was extracted using Trizol reagent (Invitrogen). Total RNA was then treated with DNase I (Invitrogen) to remove DNA and purified RNA was recovered with RNeasy® MinElute™ Cleanup kit (QIAGEN) according to the manufacturer’s instructions. Library construction and RNA sequencing were performed in Majorbio company (Shanghai, China). Briefly, RNAseq transcriptome library was prepared following TruSeq™ RNA sample preparation Kit from Illumina (San Diego, CA) using 1 μg of total RNA. Messenger RNA was isolated using oligo (dT) beads and then fragmented by fragmentation. Then, double-stranded cDNA was synthesized using a SuperScript double-stranded cDNA synthesis kit (Invitrogen, CA) with random hexamer primers (Illumina). The synthesized cDNA was then subjected to end-repair, phosphorylation and ‘A’ base addition. After size selection, paired-end RNAseq sequencing library was sequenced with the Illumina HiSeq x Ten/NovaSeq 6000 sequencer.

### Stomatal aperture measurement

Four-week-old plants were used for stomatal measurement. Stomatal images before or after high humidity treatment were collected on confocal microscope (SP8 STED, Leica) with 405 nm excitation and 435-485nm emission (to observe auto-fluorescence from the inner cell wall of the guard cell). The width and length of stomata were measured by ImageJ and the ratio of width/length was calculated. At least 50 stomatal images were analyzed for each treatment.

### Chromatin immunoprecipitation-seq (ChIPseq) and ChIP-qPCR

ChIP experiment were carried out as described previously (Liu et al., 2019). Briefly, a total of 5 g plant materials (4-weeks-old leaves of *clf-28* and Col-0 plants) grown in soil were collected and vacuum infiltrated with 1% formaldehyde solution. The leaves were then washed several times with deionized water and snap frozen in liquid nitrogen. Tissues were ground into fine powder. Chromatin was isolated and fragmented into approximately 500-1000 bp size with sonicator. The immunoprecipitation was performed with the antibody against H3 tri-methyl-Lys 27 (Upstate, USA; Cat. # 07-449). At least 2 ng of ChIP-DNA were used for Illumina library preparation according to the manufacturer’s instructions (Illumina). Library construction and sequencing (150 bp of paired-end sequences) were performed by Novogene Co. Ltd (Shanghai, China), using Illumina HiSeq 2500. Primers for ChIP-qPCR analysis are listed in Supplemental Table S3.

### ChIPseq and RNAseq analysis

Trimmomatic (version 0.36) was used to trim adaptor (Bolger et al., 2014). Next, the program Sickle (version 1.33) was used to remove bases with low quality scores (<20) and reads shorter than 20bp were eliminated. The cleaned reads were mapped to the Arabidopsis genome (TAIR10) using the Burrows–Wheeler Aligner (version 0.7.5a-r405) (Li and Durbin, 2010) for the ChIP-sequencing data and HISAT2 (version 2.1.0) (Kim et al., 2015) for mapping the RNA sequencing data, both with default settings.

Uniquely mapped reads with MAPQ>20 was collected for further analysis. The MACS (version 1.3.7) program (Zhang et al., 2008) as used to identify the read-enriched regions (peaks) of the ChIPseq data based on the following criteria: P < 1e - 5 and fold-change > 32. To quantify gene expression levels, the featureCount program of the Subread package (version 1.6.5) (Liao et al., 2013) with parameters “-s 2 -p -t exon” was used to determine the RNAseq read density. To compare expression levels across samples and genes, the RNAseq read density of each gene was normalized based on the exon length in the gene and the sequencing depth. To quantify histone markers across genes for the figure prepared with Integrative Genomics Viewer (Robinson et al., 2011), the number of reads at each position was normalized against the total number of reads (reads per million mapped reads). The DEseq program (Robinson et al., 2010) was used for detecting differentially expressed genes based on the following criteria: |log2 fold-change| > 1 and P < 0.05. The MAnorm package (Shao et al., 2012) was used for the quantitative comparison of ChIPseq signals between samples with the following criteria: |M value| > 0.5 and P < 0.05.

### Plant SA and ABA measurement

Four-weeks-old plant leaves were used for ABA and SA measurement as described previously (Huot et al., 2017). For each biological replicate, about 0.15 g leaf tissue was collected and ground into fine power in liquid nitrogen. ABA and SA were extracted using extraction buffer (20% HPLC water and 1% formic acid in methanol) at 4°C for 4-6 h, and the extracts were then air-dried in a speed vacuum and resuspended in extraction buffer. ABA and SA concentration were determined by AB SCIEX 4000Q TARP LC/MX/MX system (SCIEX QTRAP 6500+) at the mass spectrometry facility at Chinese Academy of Sciences, Center for Excellence in Molecular Plant Sciences, Shanghai.

### MAPK kinase activity assay

Four-week-old plant leaves were infiltrated with flg22 (100 nM), and leaves were harvested at indicated time points. Total proteins were extracted in protein extraction buffer (50mM Tris-HCl pH 7.5, 150mM NaCl, 5mM EDTA pH 7.5, 1mM DTT, 1% Triton X-100, 1mM Phenylmethylsulfonyl fluoride) supplemented with 1 x plant protease inhibitor cocktail (Complete EDTA-free, Roche) and 1 x phosphatase inhibitor cocktail (PhosSTOP, Roche). Bradford protein assay kit (Bio-rad) was used to determine protein concentration. An equal amount of protein was loaded onto 10% SDS-PAGE gel for western blot. Phosphorylated MPK3 and MPK6 were detected by anti-Phospho-p44/42 antibody (Cell Signaling Technology).

### ROS detection Assay

ROS measurement with luminol/HRP-based approach was performed as previously described (Yuan et al., 2021). Briefly, leaf discs of four-week-old Col-0 and *clf* plants were harvested and floated on 200 μL sterile water in a 96-well plate, and kept at room temperature under continuous light. The next morning (about 10 h later), water was replaced by a solution containing 30mg/L (w/v) luminol (Sigma-Aldrich), 20mg/L (w/v) peroxidase from horseradish (Sigma-Aldrich) and 100nM flg22. The luminescence was detected for 40 min with 1 min interval using Varioskan Flash plate reader (Thermo Fisher Scientific).

### Bacterial growth assay

*Pseudomonas syringae* pv. *tomato* DC3000 and mutant strains were cultured overnight at 30°C in Luria–Marine (LM) medium supplemented with 50 μg/mL rifampicin. The bacteria were collected by centrifugation, washed twice with water, and re-suspended in sterile water. Cell density was adjusted to OD_600_ = 0.002 (approximately 1 x 10^6^ cfu ml^-1^). Bacteria were infiltrated into leaves with a needless-syringe, and inoculated plants were kept under ambient humidity for 1-2 h to allow evaporation of excess water from the leaf. Plants were then placed back to the growth room (covered with a clear plastic dome) for disease to develop. For quantification of bacteria, 5 leaf discs from two different leaves (after surface sterilization) were taken using a cork borer (8 mm in diameter) as one biological repeat, and 3-4 repeats were taken for each treatment. Leaf discs were ground in sterile water, and bacteria solutions were diluted and plated on LM agar plates supplemented with rifampicin (at 50 mg/L). Colonies were counted with a stereoscope 24 h after incubation at 30°C.

## Data availability

Raw sequencing data from RNAseq and ChIPseq experiments were uploaded onto the Gene Expression Omnibus (GEO) database on NCBI, with the accession number GSE183559.

## Acknowledgements

The *CLF* T-DNA insertion line (SALK_139371) was kindly provided by Prof. Yue Zhou’s lab at Peking University. The *nac019/055/072* mutant seeds were kindly provided by Prof. Kenichi Tsuda’s lab at Huazhong Agricultural University. We thank the Confocal Microscopy and Imaging Facility and Metabolomics/Mass Spectrometry Facility at CAS Center for Excellence in Molecular Plant Sciences/Institute of Plant Physiology and Ecology, Shanghai, for technical support in stomatal measurement and phytohormone quantification. We thank Dr. Sheng Yang He’s lab at Michigan State University, USA, for the help with generation of the EMS screen population. This work was supported by the Chinese Academy of Sciences, Center for Excellence in Molecular Plant Sciences/Institute of Plant Physiology and Ecology, National Key Laboratory of Plant Molecular Genetics and Chinese Academy of Sciences Strategic Priority Research Program (Type-B; project number: XDB27040211).

## Author Contributions

J.W. and X-F.X. conceptualized the project and designed experiments. J.W. performed most experiments including genetic screen, water soaking assays, RNAseq, qRT-PCR, ChIP-qPCR, hormone measurement, stomatal measurement and bacterial inoculation assays. X.M. performed qRT-PCR, stomatal measurement and bacterial infection assays. J.Z. performed ChIPseq analysis. L.Y. performed ChIPseq experiment. T.C. helped with the generation of EMS population. Y.W. confirmed SNP alleles of CLF in other *sws1* lines. Y.Z. supervised ChIPseq experiment and data analysis. J.W. and X-F.X. wrote the paper with input from all authors.

---

**Supplemental Figure S1.**
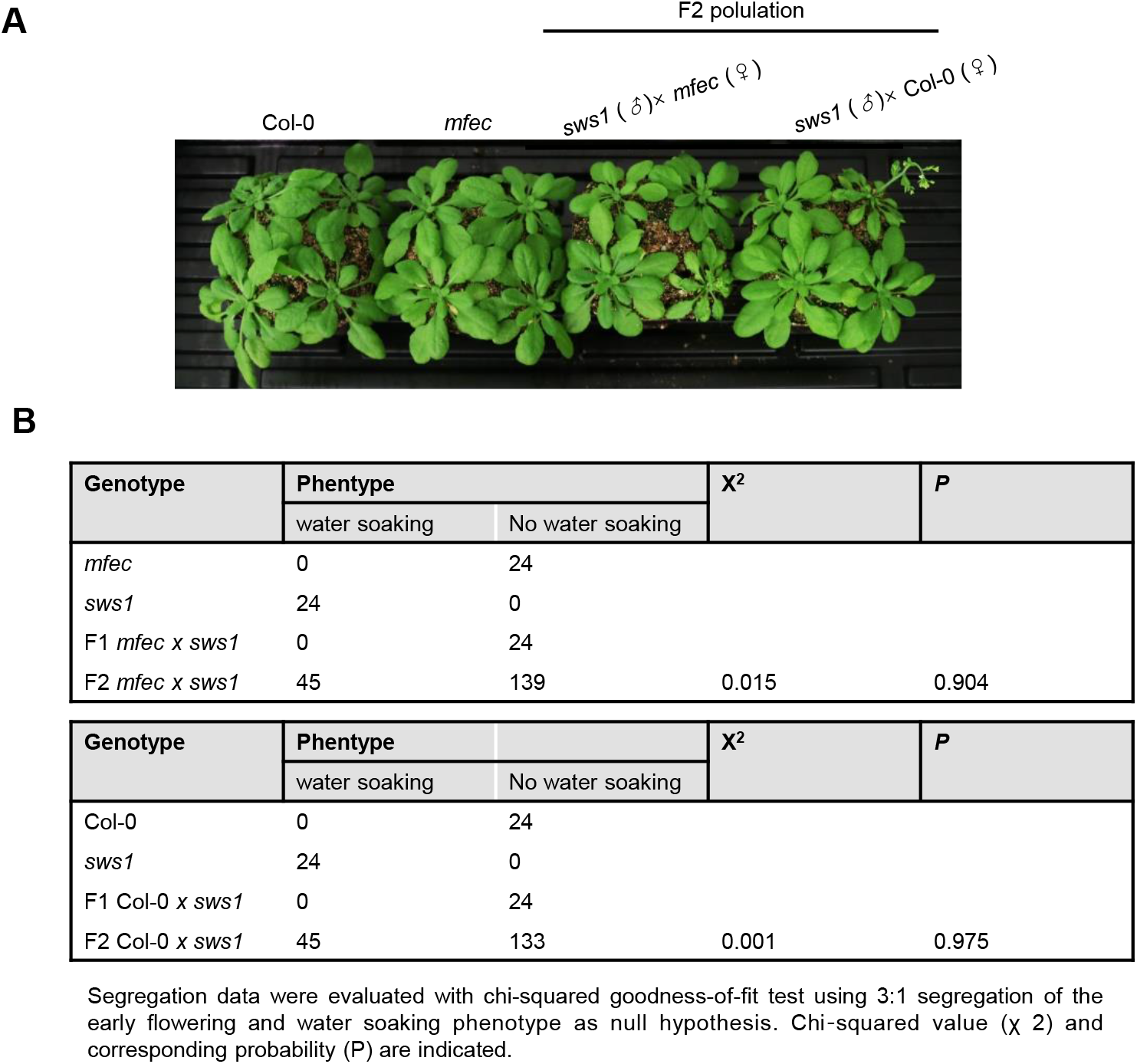
Segregation of *SWS1* allele in Col-0 or *mfec* background. (A) Morphology of 4-weeks-old Col-0, *mfec* and F2 populations of *sws1* crossed to Col-0 or *mfec* plants. Plants were grown under normal humidity (~60% relative humidity). (B) Segregation ratio of the water soaking phenotype in F1 and F2 populations of *sws1* crossed to Col-0 or *mfec* plants.

**Supplemental Figure S2.**
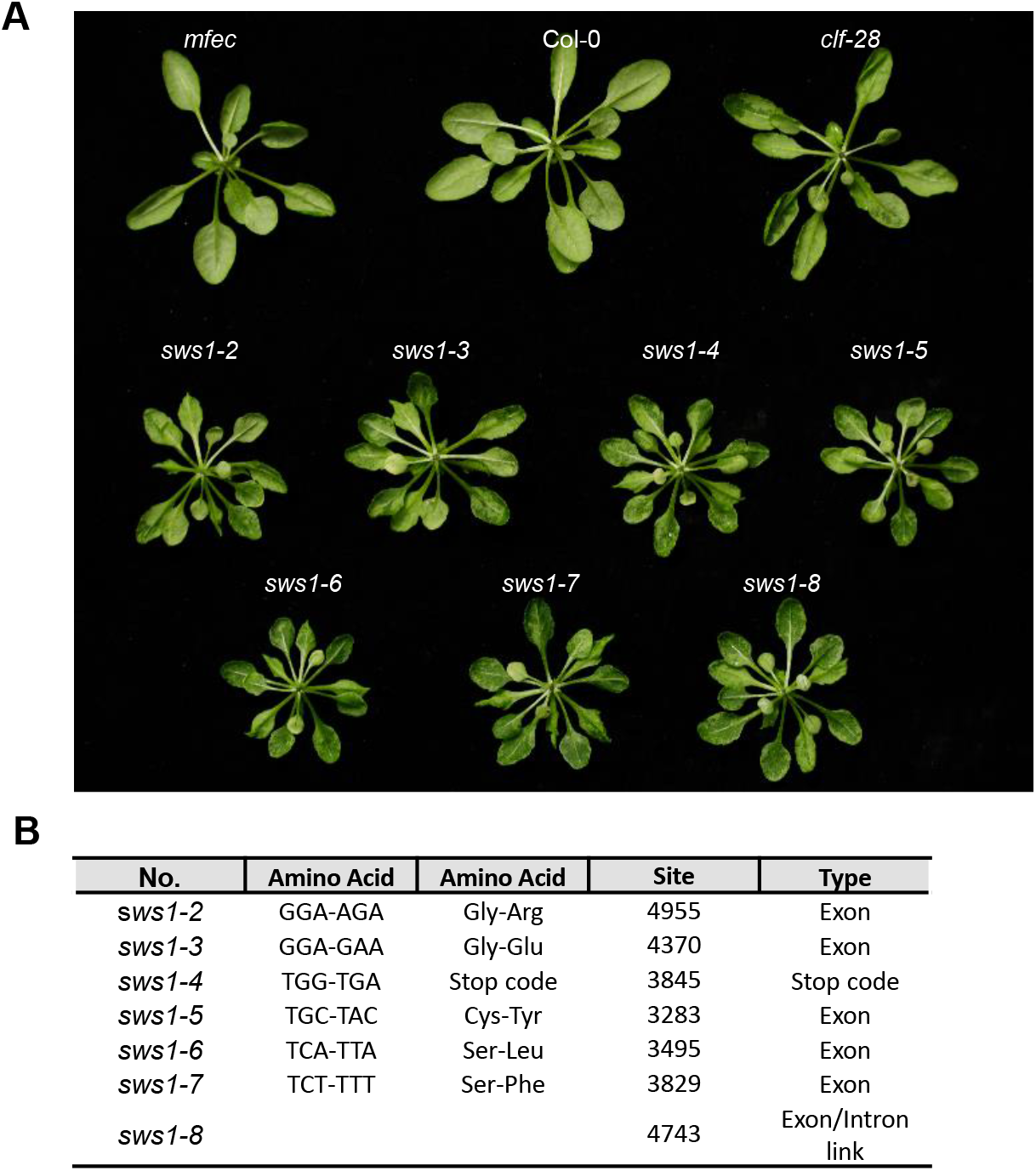
Water soaking phenotype and mutation sites of different SNP alleles of *sws1*. (A) High humidity-induced water soaking in other *sws1* lines isolated from the genetic screen. Plants were grown until 3-4 week old and were treated with high humidity for 4 h before pictures were taken. (B) Mutation type in the *CLF* gene in other *sws1* lines determined by genome sequencing.

**Supplemental Figure S3.**
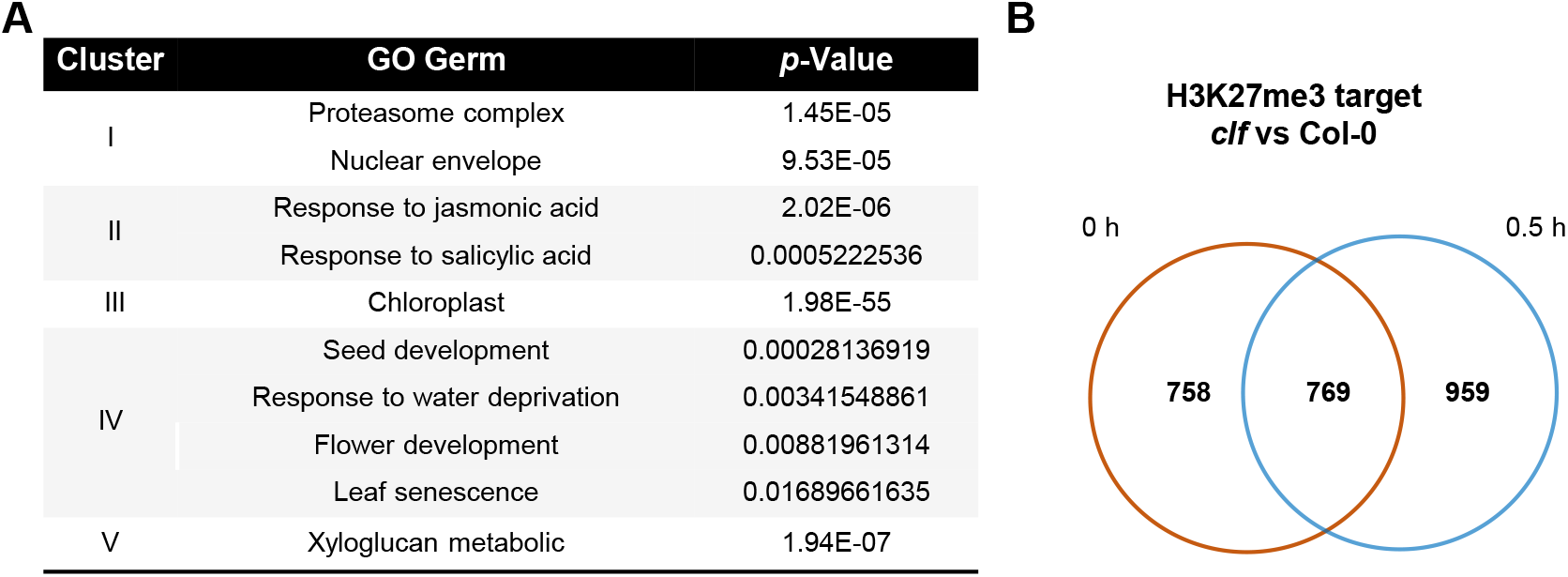
Data analysis of the RNAseq and ChIPseq results. (A) GO analysis of DEGs in each cluster as shown in Figure 4A. (B) Number of genes that show differential H3K27me3 level in the *clf* mutant compared to Col-0 plant in the ChIPseq experiment. 0 and 0.5h indicate the time of high humidity treatment.

**Supplemental Figure S4.**
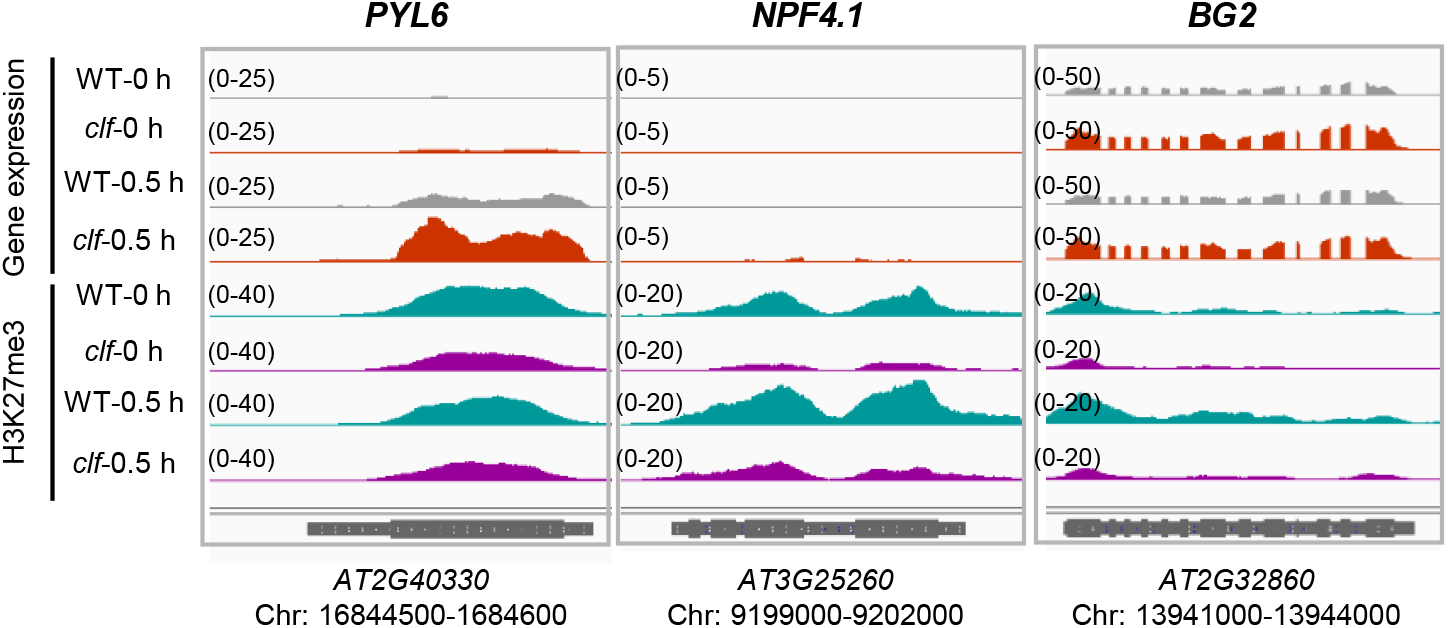
Gene expression and H3K27me3 levels of several ABA-related genes from the RNAseq and ChIPseq experiments. *PYL6, NPF4.1* and *BG2* genes show lower H3K27me3 level, accompanied with higher transcription level, in the *clf* mutant compared to Col-0 plant. 0 and 0.5 h indicate the time of high humidity treatment.

**Supplemental Figure S5.**
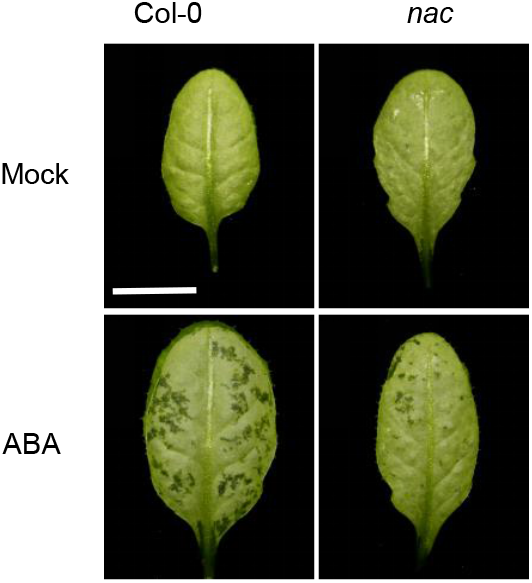
ABA-induced water soaking is significantly reduced in the *nac* triple mutant plant. Four-week-old Col-0 and *nac* plants were sprayed with ABA (50 μM) and pictures were taken 6 h later. Bar=1 cm.

## References

Blazewicz, S.J., Schwartz, E., and Firestone, M.K. (2014). Growth and death of bacteria and fungi underlie rainfall-induced carbon dioxide pulses from seasonally dried soil. Ecology 95: 1162–1172.

Bolger, A.M., Lohse, M., and Usadel, B. (2014). Trimmomatic: a flexible trimmer for Illumina sequence data. Bioinformatics 30: 2114–2120.

Bouveret, R., Schönrock, N., Gruissem, W., and Hennig, L. (2006). Regulation of flowering time by Arabidopsis MSI1. Development 133: 1693–1702.

Bratzel, F., López-Torrejón, G., Koch, M., Del Pozo, J.C., and Calonje, M. (2010). Keeping cell identity in Arabidopsis requires PRC1 RING-finger homologs that catalyze H2A monoubiquitination. Curr. Biol. 20: 1853–1859.

Chen, K., Li, G.-J., Bressan, R.A., Song, C.-P., Zhu, J.-K., and Zhao, Y. (2020). Abscisic acid dynamics, signaling, and functions in plants. J. Integr. Plant Biol. 62: 25–54.

Cutler, S.R., Rodriguez, P.L., Finkelstein, R.R., and Abrams, S.R. (2010). Abscisic acid: emergence of a core signaling network. Annu. Rev. Plant Biol. 61: 651–679.

DeFalco, T.A. and Zipfel, C. (2021). Molecular mechanisms of early plant pattern-triggered immune signaling. Mol. Cell 81: 3449–3467.

Dinneny, J.R. (2019). Developmental Responses to Water and Salinity in Root Systems. Annu. Rev. Cell Dev. Biol. 35: 239–257.

Fàbregas, N., Yoshida, T., and Fernie, A.R. (2020). Role of Raf-like kinases in SnRK2 activation and osmotic stress response in plants. Nat. Commun. 11: 6184.

Felix, G., Duran, J.D., Volko, S., and Boller, T. (1999). Plants have a sensitive perception system for the most conserved domain of bacterial flagellin. Plant J. 18: 265–276.

Fujii, H. and Zhu, J.-K. (2009). Arabidopsis mutant deficient in 3 abscisic acid-activated protein kinases reveals critical roles in growth, reproduction, and stress. Proc. Natl. Acad. Sci. U. S. A. 106: 8380–8385.

Gómez-Gómez, L. and Boller, T. (2000). FLS2: an LRR receptor-like kinase involved in the perception of the bacterial elicitor flagellin in Arabidopsis. Mol. Cell 5: 1003–1011.

González-Guzmán, M., Apostolova, N., Bellés, J.M., Barrero, J.M., Piqueras, P., Ponce, M.R., Micol, J.L., Serrano, R., and Rodríguez, P.L. (2002). The short-chain alcohol dehydrogenase ABA2 catalyzes the conversion of xanthoxin to abscisic aldehyde. Plant Cell 14: 1833–1846.

Goodrich, J., Puangsomlee, P., Martin, M., Long, D., Meyerowitz, E.M., and Coupland, G. (1997). A Polycomb-group gene regulates homeotic gene expression in Arabidopsis. Nature 386: 44–51.

Guzel Deger, A., Scherzer, S., Nuhkat, M., Kedzierska, J., Kollist, H., Brosché, M., Unyayar, S., Boudsocq, M., Hedrich, R., and Roelfsema, M.R.G. (2015). Guard cell SLAC1-type anion channels mediate flagellin-induced stomatal closure. New Phytol. 208: 162–173.

Hedrich, R. and Geiger, D. (2017). Biology of SLAC1-type anion channels - from nutrient uptake to stomatal closure. New Phytol. 216: 46–61.

Hewage, K.A.H., Yang, J.-F., Wang, D., Hao, G.-F., Yang, G.-F., and Zhu, J.-K. (2020). Chemical Manipulation of Abscisic Acid Signaling: A New Approach to Abiotic and Biotic Stress Management in Agriculture. Adv. Sci. (Weinheim, Baden-Wurttemberg, Ger. 7: 2001265.

Hsu, P.-K., Dubeaux, G., Takahashi, Y., and Schroeder, J.I. (2021). Signaling mechanisms in abscisic acid-mediated stomatal closure. Plant J. 105: 307–321.

Hu, Y., Ding, Y., Cai, B., Qin, X., Wu, J., Yuan, M., Wan, S., Zhao, Y., and Xin, X.-F. (2022). Bacterial effectors manipulate plant abscisic acid signaling for creation of an aqueous apoplast. Cell Host Microbe 30: 518–529.e6.

Huot, B., Castroverde, C.D.M., Velásquez, A.C., Hubbard, E., Pulman, J.A., Yao, J., Childs, K.L., Tsuda, K., Montgomery, B.L., and He, S.Y. (2017). Dual impact of elevated temperature on plant defence and bacterial virulence in Arabidopsis. Nat. Commun. 8: 1808.

Jensen, M. K., Kjaersgaard, T., Nielsen, M. M., Galberg, P., Petersen, K., O’Shea, C., Skriver, K. (2010) The *Arabidopsis thaliana* NAC transcription factor family: structure-function relationships and determinants of ANAC019 stress signaling. Biochem J. 426: 183–96.

Jonas, J.L., Buhl, D.A., and Symstad, A.J. (2015). Impacts of weather on long-term patterns of plant richness and diversity vary with location and management. Ecology 96: 2417–2432.

Kim, D., Langmead, B., and Salzberg, S.L. (2015). HISAT: a fast spliced aligner with low memory requirements. Nat. Methods 12: 357–360.

Kim, Y., Chung, Y.S., Lee, E., Tripathi, P., Heo, S., and Kim, K.-H. (2020). Root response to drought stress in rice *(Oryza sativa* L.). Int. J. Mol. Sci. 21.

Köhler, C. and Villar, C.B.R. (2008). Programming of gene expression by Polycomb group proteins. Trends Cell Biol. 18: 236–243.

Kuromori, T., Seo, M., and Shinozaki, K. (2018). ABA Transport and plant water stress responses. Trends Plant Sci. 23: 513–522.

Kwak, J.M., Mori, I.C., Pei, Z.M., Leonhardt, N., Torres, M.A., Dangl, J.L., Bloom, R.E., Bodde, S., Jones, J.D., and Schroeder, J.I. (2003). NADPH oxidase AtrbohD and AtrbohF genes function in ROS-dependent ABA signaling in Arabidopsis. EMBO J. 22: 2623–2633.

Kwon, C.S., Lee, D., Choi, G., and Chung, W.-I. (2009). Histone occupancy-dependent and-independent removal of H3K27 trimethylation at cold-responsive genes in Arabidopsis. Plant J. 60: 112–121.

Lajeunesse, G., Roussin-Léveillée, C., Boutin, S., Fortin, E., Laforest-Lapointe, I., Moffett, P. (2022) Light prevents pathogen-induced aqueous microenvironments via potentiation of salicylic acid signaling. bioRxiv. https://doi.org/10.1101/2022.08.10.503390.

Lau, J.A. and Lennon, J.T. (2012). Rapid responses of soil microorganisms improve plant fitness in novel environments. Proc. Natl. Acad. Sci. U. S. A. 109: 14058–14062.

Li, H. and Durbin, R. (2010). Fast and accurate long-read alignment with Burrows-Wheeler transform. Bioinformatics 26: 589–595.

Liao, Y., Smyth, G.K., and Shi, W. (2013). The Subread aligner: fast, accurate and scalable read mapping by seed-and-vote. Nucleic Acids Res. 41: e108.

Liu, C., Cheng, J., Zhuang, Y., Ye, L., Li, Z., Wang, Y., Qi, M., Xu, L., and Zhang, Y. (2019). Polycomb repressive complex 2 attenuates ABA-induced senescence in Arabidopsis. Plant J. 97: 368–377.

Liu, N., Fromm, M., and Avramova, Z. (2014). H3K27me3 and H3K4me3 chromatin environment at super-induced dehydration stress memory genes of *Arabidopsis thaliana.* Mol. Plant 7: 502–513.

Loreti, E., van Veen, H., and Perata, P. (2016). Plant responses to flooding stress. Curr. Opin. Plant Biol. 33: 64–71.

Mine, A., Berens, M.L., Nobori, T., Anver, S., Fukumoto, K., Winkelmuller, T.M., Takeda, A., Becker, D., and Tsuda, K. (2017). Pathogen exploitation of an abscisic acid-and jasmonate-inducible MAPK phosphatase and its interception by Arabidopsis immunity. Proc. Natl. Acad. Sci. U. S. A. 114: 7456–7461.

Moeder, W., Ung, H., Mosher, S., and Yoshioka, K. (2010). SA-ABA antagonism in defense responses. Plant Signal. Behav. 5: 1231–1233.

Peng, Y., Yang, J., Li, X., and Zhang, Y. (2021). Salicylic Acid: Biosynthesis and Signaling. Annu. Rev. Plant Biol. 72: 761–791.

Pu, L. and Sung, Z.R. (2015). PcG and trxG in plants - friends or foes. Trends Genet. 31: 252–262.

Ramachandran, P., Augstein, F., Nguyen, V., and Carlsbecker, A. (2020). Coping With Water Limitation: Hormones That Modify Plant Root Xylem Development. Front. Plant Sci. 11: 570.

Robinson, J.T., Thorvaldsdóttir, H., Winckler, W., Guttman, M., Lander, E.S., Getz, G., and Mesirov, J.P. (2011). Integrative genomics viewer. Nat. Biotechnol. 29: 24–26.

Robinson, M.D., McCarthy, D.J., and Smyth, G.K. (2010). edgeR: a Bioconductor package for differential expression analysis of digital gene expression data. Bioinformatics 26: 139–140.

Roussin-Léveillée, C., Lajeunesse, G., St-Amand, M., Veerapen, V.P., Silva-Martins, G., Nomura, K., Brassard, S., Bolaji, A., He, S.Y., and Moffett, P. (2022). Evolutionarily conserved bacterial effectors hijack abscisic acid signaling to induce an aqueous environment in the apoplast. Cell Host Microbe 30: 489–501.e4.

Schwartz, A.R., Morbitzer, R., Lahaye, T., and Staskawicz, B.J. (2017a). TALE-induced bHLH transcription factors that activate a pectate lyase contribute to water soaking in bacterial spot of tomato. Proc. Natl. Acad. Sci. U. S. A. 114: E897–E903.

Schwartz, A.R., Morbitzer, R., Lahaye, T., and Staskawicz, B.J. (2017b). TALE-induced bHLH transcription factors that activate a pectate lyase contribute to water soaking in bacterial spot of tomato. Proc. Natl. Acad. Sci. U. S. A. 114: E897–E903.

Shao, Z., Zhang, Y., Yuan, G.-C., Orkin, S.H., and Waxman, D.J. (2012). MAnorm: a robust model for quantitative comparison of ChIP-Seq data sets. Genome Biol. 13: R16.

Siao, W., Coskun, D., Baluška, F., Kronzucker, H.J., and Xu, W. (2020). Root-Apex Proton Fluxes at the Centre of Soil-Stress Acclimation. Trends Plant Sci. 25: 794–804.

Singkaravanit-Ogawa, S., Kosaka, A., Kitakura, S., Uchida, K., Nishiuchi, T., Ono, E., Fukunaga, S., and Takano, Y. (2021). Arabidopsis CURLY LEAF functions in leaf immunity against fungal pathogens by concomitantly repressing SEPALLATA3 and activating ORA59. Plant J. 108: 1005–1019.

Takagi, H. et al. (2013). QTL-seq: rapid mapping of quantitative trait loci in rice by whole genome resequencing of DNA from two bulked populations. Plant J. 74: 174–183.

Takasaki, H., Maruyama, K., Takahashi, F., Fujita, M., Yoshida, T., Nakashima, K., Myouga, F., Toyooka, K., Yamaguchi-Shinozaki, K., and Shinozaki, K. (2015). SNAC-As, stress-responsive NAC transcription factors, mediate ABA-inducible leaf senescence. Plant J. 84: 1114–1123.

Taketani, R.G., Lançoni, M.D., Kavamura, V.N., Durrer, A., Andreote, F.D., and Melo, I.S. (2017). Dry Season Constrains Bacterial Phylogenetic Diversity in a Semi-Arid Rhizosphere System. Microb. Ecol. 73: 153–161.

Tran, L.S., Nakashima, K., Sakuma, Y., Simpson, S.D., Fujita, Y., Maruyama, K., Fujita, M., Seki, M., Shinozaki, K., Yamaguchi-Shinozaki, K. (2004) Isolation and functional analysis of Arabidopsis stress-inducible NAC transcription factors that bind to a drought-responsive cis-element in the early responsive to dehydration stress 1 promoter. Plant Cell. 16: 2481–98.

Turck, F., Roudier, F., Farrona, S., Martin-Magniette, M.-L., Guillaume, E., Buisine, N., Gagnot, S., Martienssen, R.A., Coupland, G., and Colot, V. (2007). Arabidopsis TFL2/LHP1 specifically associates with genes marked by trimethylation of histone H3 lysine 27. PLoS Genet. 3: e86.

Wildermuth, M.C., Dewdney, J., Wu, G., and Ausubel, F.M. (2001). Isochorismate synthase is required to synthesize salicylic acid for plant defence. Nature 414: 562–565.

Xiao, J. and Wagner, D. (2015). Polycomb repression in the regulation of growth and development in Arabidopsis. Curr. Opin. Plant Biol. 23: 15–24.

Xin, X.-F., Nomura, K., Aung, K., Velásquez, A.C., Yao, J., Boutrot, F., Chang, J.H., Zipfel, C., and He, S.Y. (2016). Bacteria establish an aqueous living space in plants crucial for virulence. Nature 539: 524–529.

Xu, L. and Shen, W.H. (2008). Polycomb silencing of KNOX genes confines shoot stem cell niches in Arabidopsis. Curr Biol 18: 1966–1971.

Yoshida, R., Umezawa, T., Mizoguchi, T., Takahashi, S., Takahashi, F., and Shinozaki, K. (2006). The regulatory domain of SRK2E/OST1/SnRK2.6 interacts with ABI1 and integrates abscisic acid (ABA) and osmotic stress signals controlling stomatal closure in Arabidopsis. J. Biol. Chem. 281: 5310–5318.

Zhang, A., Ren, H.-M., Tan, Y.-Q., Qi, G.-N., Yao, F.-Y., Wu, G.-L., Yang, L.-W., Hussain, J., Sun, S.-J., and Wang, Y.-F. (2016). S-type Anion Channels SLAC1 and SLAH3 Function as Essential Negative Regulators of Inward K+ Channels and Stomatal Opening in Arabidopsis. Plant Cell 28: 949–955.

Zhang, X., Germann, S., Blus, B.J., Khorasanizadeh, S., Gaudin, V., and Jacobsen, S.E. (2007). The Arabidopsis LHP1 protein colocalizes with histone H3 Lys27 trimethylation. Nat. Struct. Mol. Biol. 14: 869–871.

Zhang, Y., Liu, T., Meyer, C.A., Eeckhoute, J., Johnson, D.S., Bernstein, B.E., Nusbaum, C., Myers, R.M., Brown, M., Li, W., and Liu, X.S. (2008). Model-based analysis of ChIP-Seq (MACS). Genome Biol. 9: R137.

Zhao, Y. et al. (2018). Arabidopsis Duodecuple Mutant of PYL ABA Receptors Reveals PYL Repression of ABA-Independent SnRK2 Activity. Cell Rep. 23: 3340–3351 e5.

Zhao, Y., Gao, J., Im Kim, J., Chen, K., Bressan, R.A., and Zhu, J.-K. (2017). Control of Plant Water Use by ABA Induction of Senescence and Dormancy: An Overlooked Lesson from Evolution. Plant Cell Physiol. 58: 1319–1327.

Zheng, X.-Y., Spivey, N.W., Zeng, W., Liu, P.-P., Fu, Z.Q., Klessig, D.F., He, S.Y., and Dong, X. (2012). Coronatine promotes *Pseudomonas syringae* virulence in plants by activating a signaling cascade that inhibits salicylic acid accumulation. Cell Host Microbe 11: 587–596.

Zhou, Y., Tergemina, E., Cui, H., Förderer, A., Hartwig, B., Velikkakam James, G., Schneeberger, K., and Turck, F. (2017). Ctf4-related protein recruits LHP1-PRC2 to maintain H3K27me3 levels in dividing cells in *Arabidopsis thaliana*. Proc. Natl. Acad. Sci. U. S. A. 114: 4833–4838.

Zhu, J.-K. (2016). Abiotic Stress Signaling and Responses in Plants. Cell 167: 313–324.

